# Affinity-based polymers provide long-term immunotherapeutic drug delivery across particle size ranges optimal for macrophage targeting

**DOI:** 10.1101/802801

**Authors:** Nathan A. Rohner, Linda Purdue, Horst A. von Recum

## Abstract

Long term drug delivery to specific arms of the immune system can be technically challenging to provide limited off-target toxicity as well as prolonged delivery and specific cellular targeting given the limits of current drug delivery systems. In this work, we demonstrate the robustness of a cyclodextrin (CD) polymer platform that can extend immunomodulatory drug delivery via affinity interactions to promote long-term, sustained release at multiple size scales. The parameter space of synthesis variables (pre-incubation and stirring speed) and post-synthesis grinding effects on resulting particle diameter were characterized. We demonstrate that polymerized CD forms exhibit size-independent release profiles of the small molecule drug lenalidomide (LND) and can provide similar drug delivery profiles as macro-scale CD polymer disks. CD polymer microparticles and nanoparticles demonstrated no significant cytotoxicity as compared to the base CD macromonomer when co-incubated with fibroblasts. Uptake of ground CD nanoparticles was significantly higher following incubation with RAW 264.7 macrophages in culture over originally synthesized, larger CD microparticles. Thus, the affinity/structure properties afforded by polymerized CD allow particle size to be modified to affect cellular uptake profiles independently of drug release rate for applications in cell-targeted drug delivery.

## 1. Introduction

Sustained, localized lenalidomide (LND) delivery is not clinically available, yet could improve patient outcomes and reduce associated costs for many chronic disease scenarios. LND is used clinically to treat multiple myeloma and chronic lymphocytic leukemia and has shown promise in pre-clinical studies for other chronic and debilitating diseases. However, the cost of frequent, often daily treatments, especially due to rising drug prices, resulted in an average charge per patient per month of $14,656 for multiple myeloma in 2014 ^1^. Another disease requiring frequent LND dosing, chronic lymphocytic leukemia has a total annual cost in the United States of $5.13 billion. The per-patient lifetime cost is $604,000, mostly due to oral targeted therapies becoming the first-line treatment ^2^. Despite the monetary costs, there are major benefits to LND therapy for multiple myeloma and chronic lymphocytic leukemia patients. Subjects with chronic lymphocytic leukemia treated with LND experience a longer time of progression-free survival (median 33.9 months) ^3^. Moreover, current clinical studies illustrate that LND can extend time to subsequent treatment without affecting the ability of a patient to respond to subsequent therapy ^4^. In the case of multiple myeloma, patients treated with LND exhibited a 15% 8-year overall survival increase in comparison to the placebo group ^5^. Adding even more potential, LND treatment of Parkinson’s and multiple sclerosis is demonstrating benefits in pre-clinical animal models. With a total cost each year of $25 billion, Parkinson’s treatment is in need of better options as for a patient in long-term care, the annual cost with psychosis is $31,178 and without psychosis is $14,461 ^6^. Multiple sclerosis costs $2.5 billion in the United States, with lifetime costs for patients exceeding $4 million ^7^.

In clinical applications, repeated oral administration of LND is needed to be effective due to a half of bioavailability of approximately 3 hours ^2^. Two separate trials were conducted to analyze the impact of treating multiple myeloma with LND. The IFM 2005-02 study ran two 28 day cycles, where subjects were given 25 mg/day of LND on days 1-21 of the cycle. The mean time of treatment was 25 months, which was then followed by maintenance treatment within the six months following autologous stem-cell transplantation. Once a day, 10 mg of LND were given to the subject for the first three months of maintenance treatment. If the subject did not have a dose-limiting toxicity, the dosage was increased to 15 mg. For the second trial, CALGB 100104, mean initial treatment time was extended to 30 months. It followed the same procedure for maintenance treatment as the IFM 2005-02 study, but maintenance was started 100-110 days after ASCT ^5^. The CONTINUUM trial was conducted to determine the effects of treating chronic lymphocytic leukemia with LND. For the first 28 day cycle, the subject received 2.5 mg of LND once a day. If this dose was well tolerated, the second 28 cycle was increased to a 5 mg dose per day. The dosage was increased to 10 mg per day only if five continuous cycles of 5 mg/day were well tolerated and they had not received a minimal residual disease negative complete response ^3–4^. LND is also required frequent dosing in animal models, exploring the possible treatment of Parkinson’s disease. The dosing schedule was found to have protective effects for amyotrophic lateral sclerosis. The solution containing LND was made fresh every week, and was administered via oral gavage five times a week in a 5 mL/kg volume ^8^. Furthermore, an animal model of multiple sclerosis (MS) exhibiting experimental autoimmune encephalomyelitis (EAE), has been treated therapeutically with LND. After the appearance of clinical symptoms, 30 mg/kg in 0.9% CMC-Na of LND was administered daily to the mice ^9^. Additional factors further hinder LND efficacy as LND has poor solubility; 82% of the administered dose is excreted unchanged in the urine ^5^ and it is unable to penetrate the blood-brain barrier (BBB) effectively, especially when compared to its parent molecule thalidomide ^8^.

LND has proven to be beneficial, but current therapy is wasteful, expensive and requires continuous dosing that relies heavily upon patient compliance and exhibits some concern for off-target effects. Oral LND doses deliver very little to the target cells as most of the dosage is excreted ^2, 5^. LND also has poor solubility, increasing the difficulty of successfully delivering it to the area of treatment. In previous work, we have leveraged polymerized CDs (pCD) to continuously deliver hydrophobic, small molecule drugs on the order of weeks to months by leveraging affinity interactions ^10–21^. A high concentration of affinity complexation sites to delay drug release is achieved by polymerizing CD monomers into larger disk structures, coatings, and more ^11–13, 22–26^. Herein, the synthesis variables for polymerizing CD microparticles and the resulting device size effects are explored for reducing CD-associated cytotoxicity, providing sustained LND delivery with a single application, and promoting biological effectiveness *in vitro*.

## 2. Materials and Methods

### 2.1 Materials

α-cyclodextrin (α-CD) prepolymer, β-cyclodextrin (β-CD) prepolymer, γ-cyclodextrin (γ-CD) prepolymer, lightly crosslinked with epichlorohydrin, and octakis (6-deoxy-6-amino) γ-CD octahydrochloride were purchased from CycloLab (Budapest, Hungary). Ethylene glycol diglycidyl ether was purchased from Polysciences, Inc. (Warrington, PA). Hexamethylene diisocyanate were purchased from Sigma-Aldrich (St. Louis, MO). All other reagents, solvents, and chemicals were purchased from Thermo Fisher Scientific (Waltham, MA). LND was purchased from Cayman Chemicals (Ann Arbor, MI).

### 2.2 Methods

#### 2.2.1 Microparticle Synthesis

The synthesis process begins with either epichlorohydrin-crosslinked α-CD, β-CD, or γ-CD prepolymyer solubilized in 0.2 M potassium hydroxide (25% w/v) and pre-heated to 60°C for 20 minutes. A 50:1 light mineral oil to Tween85/Span85 solution (24%/76%) was mixed on a stir plate at 60°C. Ethylene glycol diglycidyl ether was added drop-wise to the solution. Next, the solution was vortexed to ensure homogenous distribution of the crosslinker before being poured into the oil/Tween85/Span85 mixture. Temperature was then increased to 70°C. After three hours, the polymerized CD microparticles were formed. The microparticles were centrifuged to get rid of excess oil, washed with excess hexanes twice, and finally washed with deionized water twice. The microparticles were resuspended, frozen, and lyophilized until dry before further use. β-CD particles were used to provide consistency among methods in experiments for validating synthesis parameter effects on size, evaluating grinding time effects, and determining viability, yet similar results are achievable with the other CD prepolymers. γ-CD prepolymer (selected due to its predicted greater affinity binding (66.5±22.5 μM) with LND) was tested in LND release and macrophage uptake studies to demonstrate application.

#### 2.2.2 Polydispersity Index (PDI) Calculations

The PDI of the synthesized microparticles was calculated by dividing the standard deviation by the size average and then squaring the resulting value. The PDI expresses uniformity of the synthesized particles in solution.

#### 2.2.3 Scanning Electron Microscope Imaging

Samples were prepared by gold sputter coating after mounting to the SEM (scanning electron microscope) stage with carbon tape. Images were taken using a JSM-6510LV SEM (Peabody, MA) under high vacuum with a 15kV accelerating voltage at a working distance of 14 mm and scan time of 20 seconds. Magnification and scale bars are as indicated in the figure captions and images.

#### 2.2.4 Particle Grinding

Polymerized microparticles were transferred into a porcelain mortar. Pestle grinding in a consistent, circular motion was continued for a pre-determined time period. Deionized water was added to the mortar, and the solution pipetted out with multiple washes ensuring maximal recovery of the ground particle population.

#### 2.2.5 Disk Synthesis

To make polymer disks for comparison to γ-CD particles, vacuum dried epichlorohydrin-crosslinked γ-CD prepolymer was dissolved in dimethylsulfoxide (DMSO) as a 25% w/v solution and heated. 1,6-hexamethylene diisocyanate (HDI) was added to the solution and then vortexed. Bubbles were removed by gentle stirring. The homogenous solution was poured into a Teflon dish, covered with parafilm, and allowed to crosslink until it completely solidified. The polymer film was cut into disks using an 8mm circular biopsy punch. The resulting disks were washed in sequence with excess DMSO for 24 hours, then 50/50 DMSO and deionized water the next day, and finally deionized water alone for 3 days before drying.

#### 2.2.6 Viability and Proliferation Assay

National Institute of Health NIH/3T3 fibroblasts (ATCC.org, Manassas, VA) were cultured in Dulbecco’s Modified Eagle’s Medium (DMEM) supplemented with fetal bovine serum (FBS) and penicillin/streptomycin at 37°C with 5% CO_2_. Cells were plated at 50,000 cells per well in a 24 well tissue culture treated plate and allowed to adhere in media. Autoclave sterilized β-CD polymer microparticles, ground β-CD particles, or β-CD monomer were added either directly to the cell wells or into 0.4 micron cutoff well inserts (Thermo Fisher Scientific, Waltham, MA) at equivalent concentrations of 50 μg/mL, and incubated for 72 hours. 100 μl of 0.15 mg/ml resazurin (Thermo Fisher Scientific, Waltham, MA), was added to evaluate metabolic activity in the alamarBlue assay for quantifying fibroblast viability and proliferation ^27^, per treatment well. After incubation for an additional 24 hours, fluorescence at 530/590 nm was measured with a Synergy H1 Microplate Reader (BioTek Instruments, Inc., Winooski, VT) and results were normalized to media only controls.

#### 2.2.7 LND Drug Loading and Release

This release tested three polymer size conditions: γ-CD disks, regular sized γ-CD microparticles, and ground γ-CD microparticles. Each condition was composed of three sample aliquots, containing 20 mg of polymer within. To account for any drug remaining in the tube, a control aliquot that did not contain any CD was analyzed. A stock loading solution of LND in DMSO was made at 10 mg/mL. Each sample aliquot received 1 mL of loading solution and the CD was allowed to complex with the LND for 3 days. During the loading period and throughout the release, the samples were kept on a shaker with a regulated temperature of 37°C. After the loading period concluded, a daily sample of release media was taken with complete release buffer replacement. The aliquots were centrifuged down and the excess solution was pipetted off to ensure no removal of polymers.

#### 2.2.8 Macrophage Particle Uptake

RAW 264.7 macrophages were cultured in DMEM supplemented with FBS and penicillin/streptomycin at 37°C with 5% CO_2_. The cells were seeded onto coverslips and allowed to adhere before addition of rhodamine-labeled γ-CD microparticles and γ-CD ground particles into culture media. Samples were gently agitated to mix and cells were allowed to incubate with the particle suspensions for 72 hours. Coverslips were then washed 3x in PBS before fixing in 4% paraformaldehyde and mounted with Vectashield (Vector Laboratories, Inc., Burlingame, CA) on glass slides for imaging with a Zeiss Observer.Z1 microscope equipped with an AxioCam ERc5s (Carl Zeiss AG, Oberkochen, Germany)

## 3. Results

### 3.1 Microparticle Synthesis Setup Allows for Precise Parameter Control

The need for new multi-functional biomaterials is evident given the resistances and barriers to conventional therapeutic approaches for many diseases. The use of polysaccharide-based polymers as biomaterial depots and in functional coatings for the delivery of small molecule drugs, such as antibiotics and anti-inflammatory therapeutics has seen steady improvement ^11, 13, 18, 20, 28–29^. Previously, polymerized CDs (pCD) were leveraged to continuously deliver small molecule drugs on the order of weeks to months by leveraging affinity interactions ^10–11, 16, 19–21, 25, 28, 30–31^. However, a variety of particle sizes may be necessary for targeting different tissues and cellular subtypes ^32–33^. Current batch synthesis of CD polymer particles is accomplished via inverse emulsion polymerization (Figure 1).

**Figure 1.**
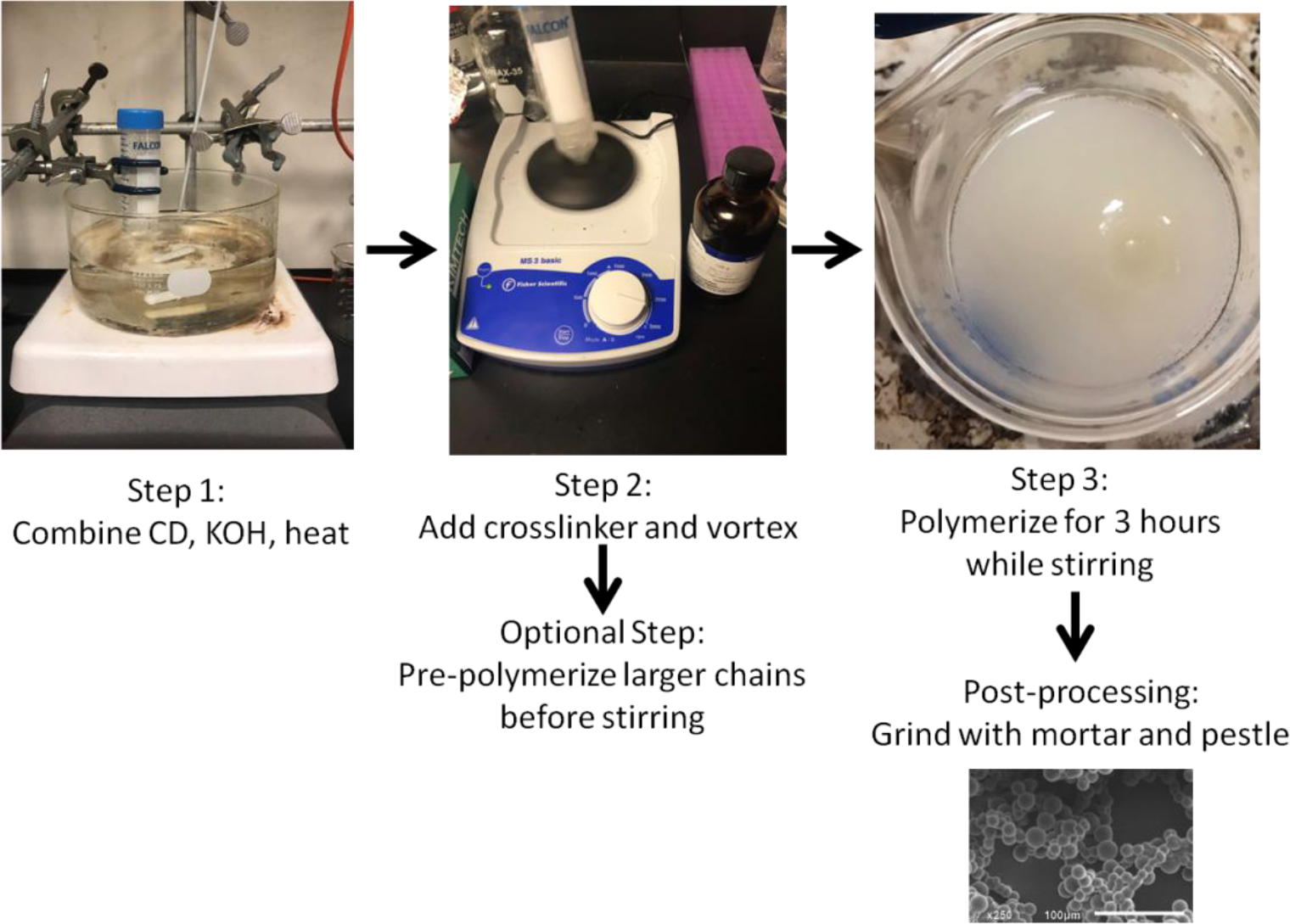
Synthesis schematic and pictures depicting process. SEM image of ground particles.

### 3.2 Stir Speed and Pre-Polymerization Time Tune Resulting β-CD Particle Size

During CD particle synthesis certain parameters were varied to determine impact on the size of the polymerized particles. One such change was the speed of the stir bar during the three hour synthesis time. The range of speeds tested were 300–1000 RPM, which still resulted in feasible particle polymerization. It was determined that increased stirring speed resulted in smaller polymerized particles (Figure 2, top). The effect of pre-polymerization before starting the reverse emulsion polymerization with stirring was also tested during microparticle synthesis. Crosslinker was added and the solution was vortexed, but was not poured into the oil/Tween85/Span85 mixture until the indicated time. The range of feasible pre-polymerization time included 0–120 minutes. It was found that as the length of pre-polymerization time increased, the size of the polymerized CD microparticles also increased (Figure 2, bottom).

**Figure 2.**
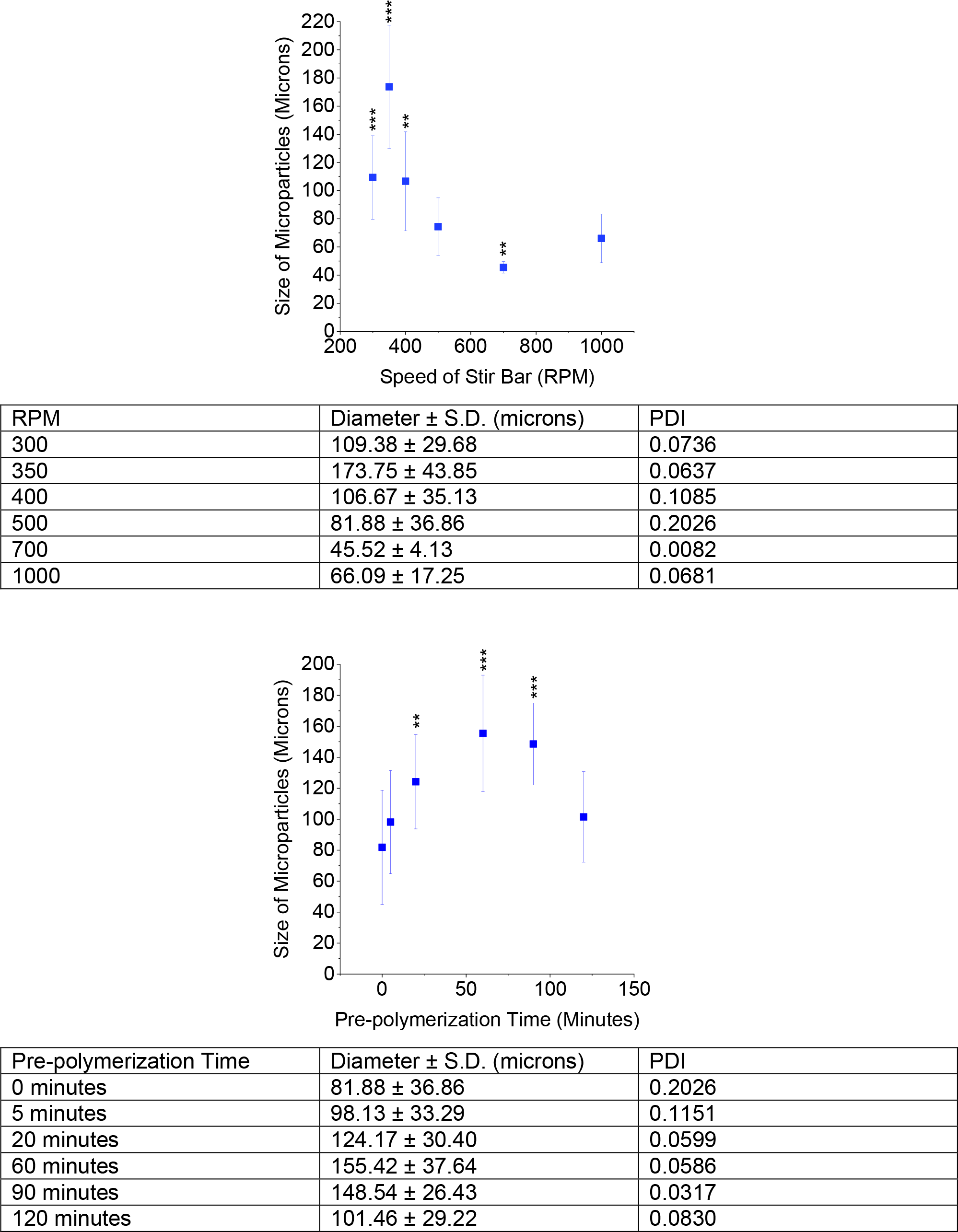
Effect of modifying RPM (top) or pre-polymerization time (bottom) on CD particle size. * indicates p < 0.05, ** p < 0.01, *** p < 0.001 by one-way ANOVA followed by Bonferroni multiple comparisons versus the 500 RPM results. n = 20 measurements from at least 2 batches per condition.

### 3.3 Versatile CD Polymer Particles Exhibit Robust Stability

The polymerized forms of alpha, beta, and gamma monomer-derivatives were compared to one another. Minor differences in size was measured between the different CD types (Figure 3, top). To test particle stability, polymerized β-CD microparticles were exposed to a range of pH between 4 and 10. It was found that despite the wide range in pH, the microparticles demonstrated no significant change in size (Figure 3, bottom). Polymerized microparticles were also sonicated, which has been traditionally used to break up particle clumps that are not thoroughly crosslinked. Tubes containing the microparticles were placed in a bath sonication machine, which applied sonic waves to the particles. Different sonication times were tested, from 1-60 minutes, but there was no significant change in microparticle size (data not shown).

**Figure 3.**
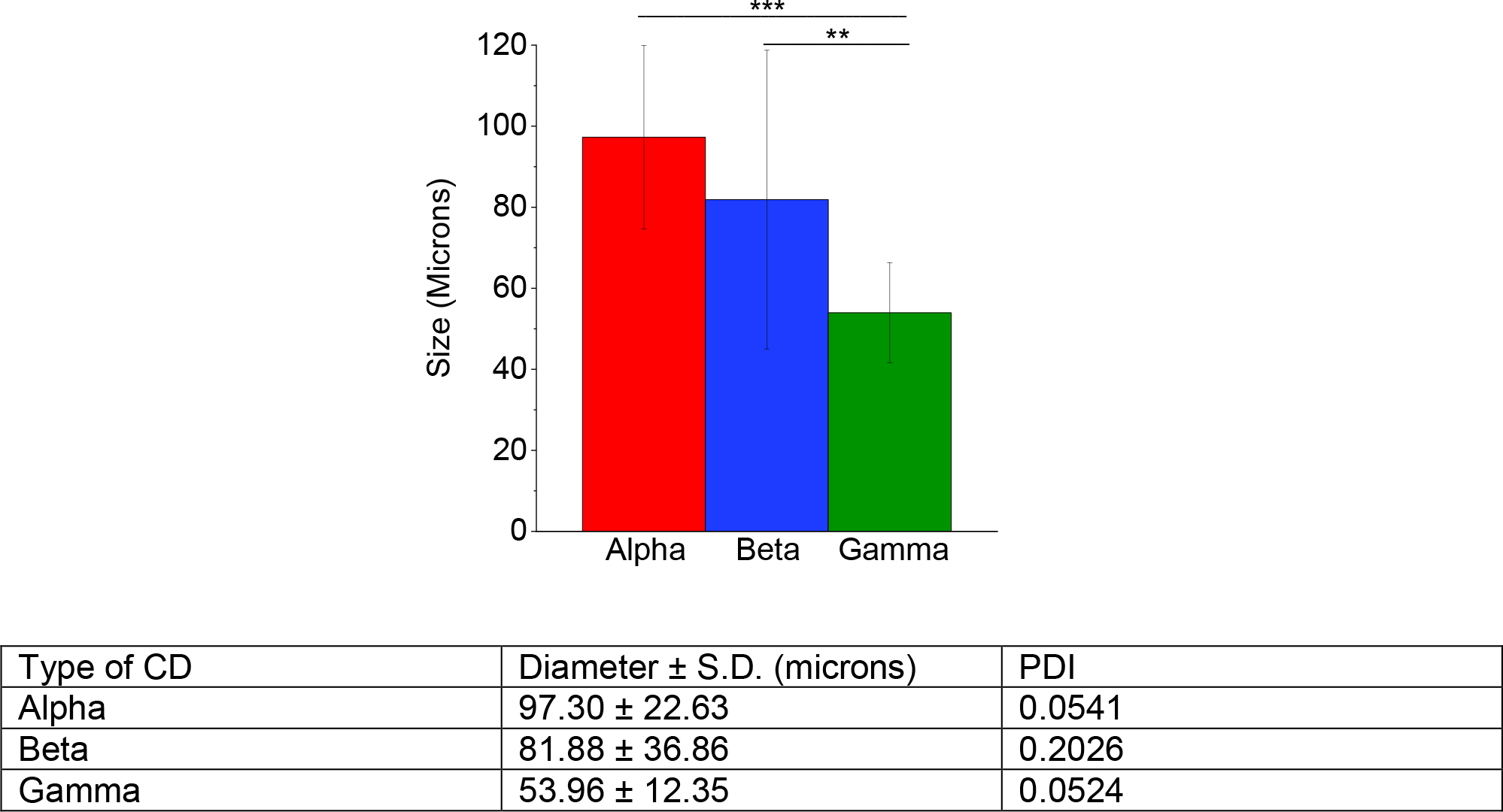
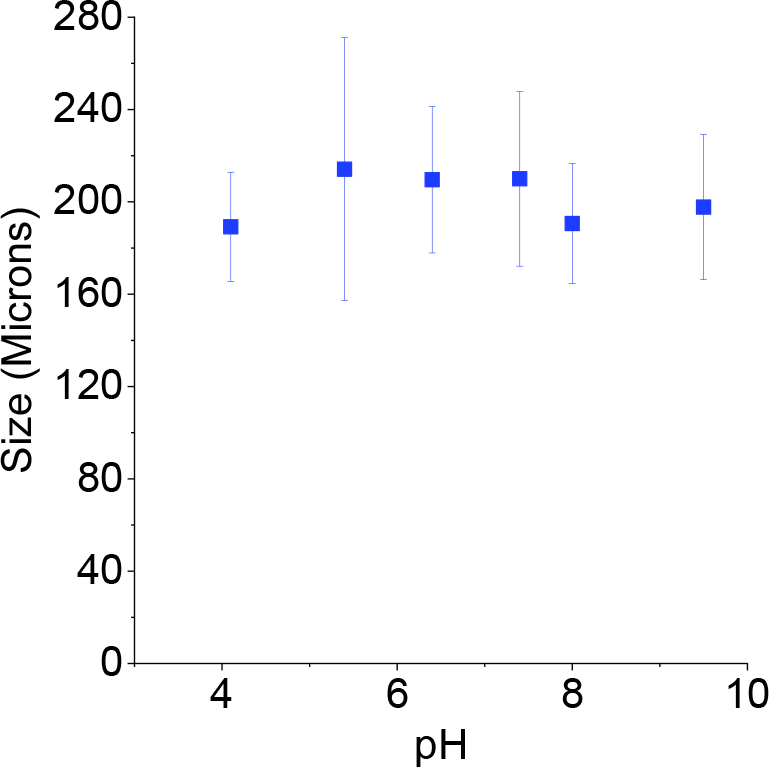
Testing polymeric CD particle variations and robustness, we observe only minor changes in size when synthesized from various CD pre-polymer types (top). Example β-CD particle size does not change significantly with change in pH from 4-10 (bottom). * p < 0.05 by one-way ANOVA with Tukey post-hoc (top). n = 20 measurements from at least 2 batches per condition.

### 3.4 Polymeric β-CD Particle Size Modifications Post-Synthesis

Traditional particles, liposomes, and shell-core drug delivery systems require changes in synthesis to modify physical parameters and are even then limited in size modifications. However, it was hypothesized that polymerized CD would be amenable to post-processing techniques due to the polymeric crosslinking structure while still maintaining effective release profiles via their affinity-based release mechanism. To determine if microparticle size could be changed post-synthesis, microparticles were ground for a pre-determined amount of time using a mortar and pestle. They were ground for either one, five, or ten minutes. Samples of each time point were taken, and it was determined that while one minute did not significantly reduce pCD particle diameters, five and ten minutes reduced the diameters to 61% and 36% of original size, respectively (Figure 4, left). Moreover, SEM images of ground particles depicted an overall spherical shape similar to the original synthesized CD particles (Figure 4, right).

**Figure 4.**
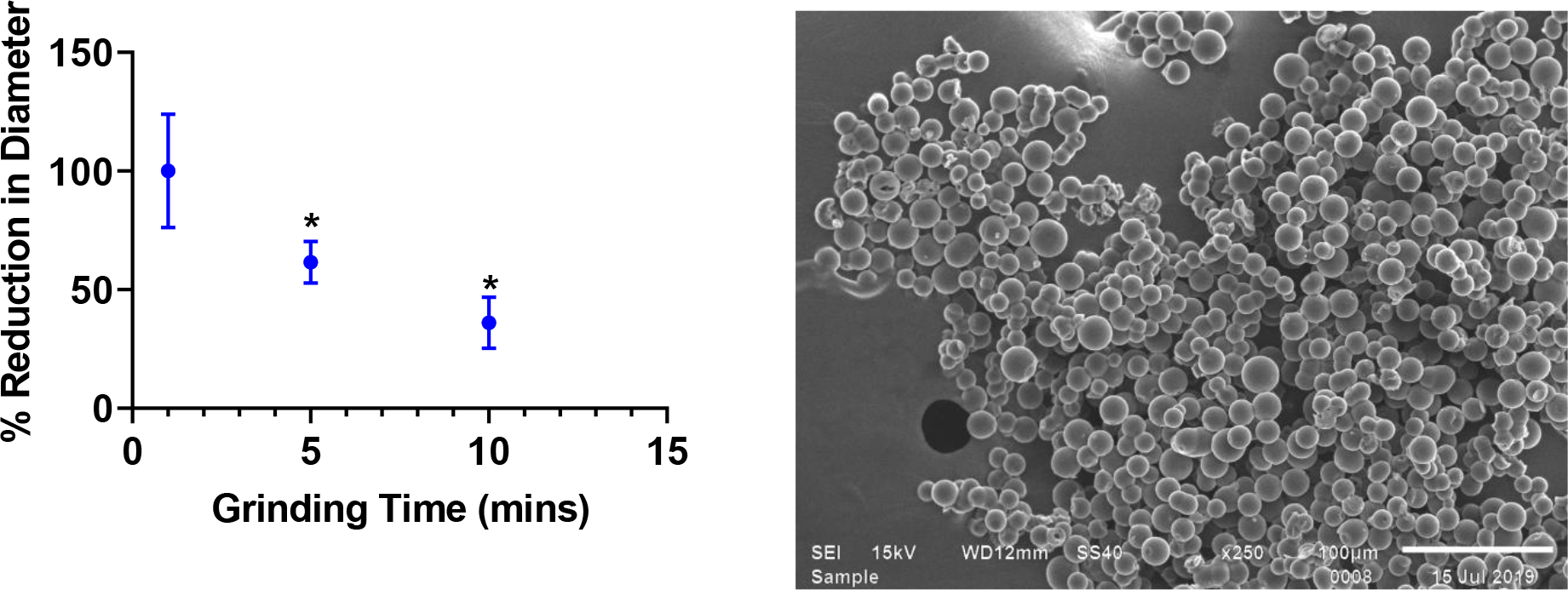
Polymeric CD particle diameter changes upon grinding time with mortar and pestle with measurements of particles for samples at different time points. SEM images of dry, ground polymerized CD particles indicate surfaces are still geometrically spherical even after grinding. * indicates p < 0.05 by one-way ANOVA with Tukey post-hoc. n = 20 measurements per condition.

### 3.5 β-CD Polymer Particles Exhibit No Adverse Cytotoxicity as Compared to Monomer

Previous studies have indicated that CD monomer has a potential cytotoxic effect, sequestering cholesterol and other molecules from the body. Polymerized CD microparticles and smaller ground particles were co-incubated with NIH/3T3 cells for 24 hours and proliferation and viability was determined relative to media only control wells. While pCD-β-MP and ground pCD-β-MP exhibited excellent cytocompatibility whether in direct incubation or separated from cells via a transwell (porous tissue culture dish insert), the β-CD monomer was observed to reduce proliferation of cells by almost 60% after 24 hours incubation (Figure 5).

**Figure 5.**
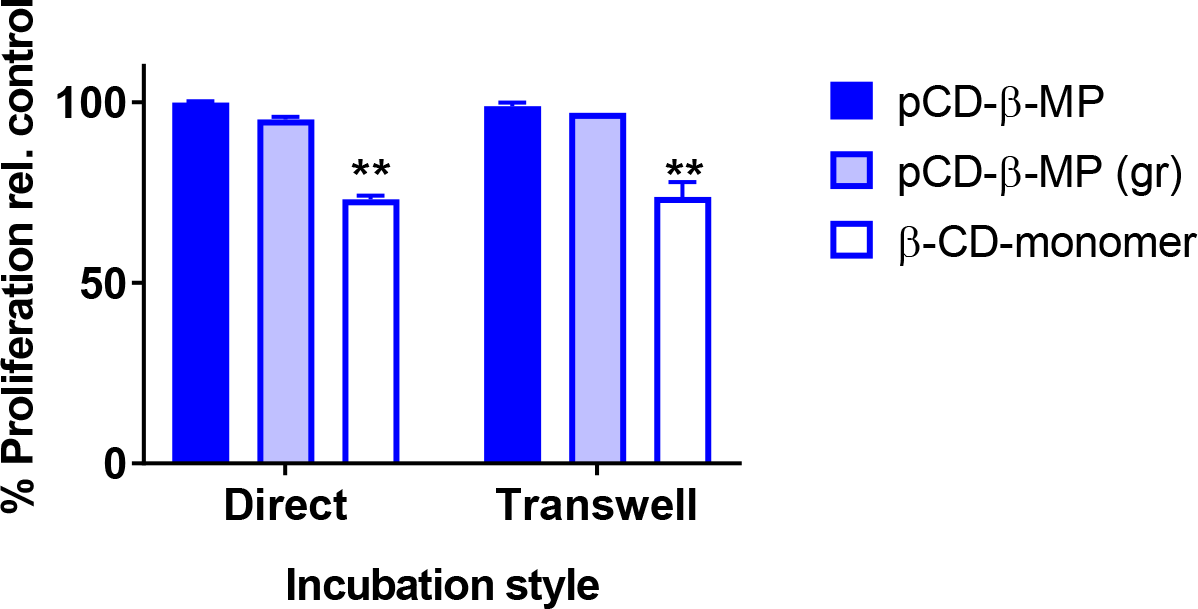
Direct and indirect CD cytotoxicity to NIH/3T3 cells in co-culture with microparticles, ground particles, and CD monomer relative to untreated controls. ** indicates p < 0.01 by two-way ANOVA with Tukey test post-hoc versus the healthy, normal growth of the control group. n=3 replicates per condition.

### 3.6 γ-CD Polymer Macrostructures Extend LND Drug Release

Synthesized CD particles were prepared and loaded with LND for 72 hours followed by release into media. Daily release aliquots were measured with complete replacement of media. While pCD disks exhibited significant LND drug release for on average 7 days, the pCD-γ-MP and ground particle forms released for up to 10-13 days.

**Figure 6.**
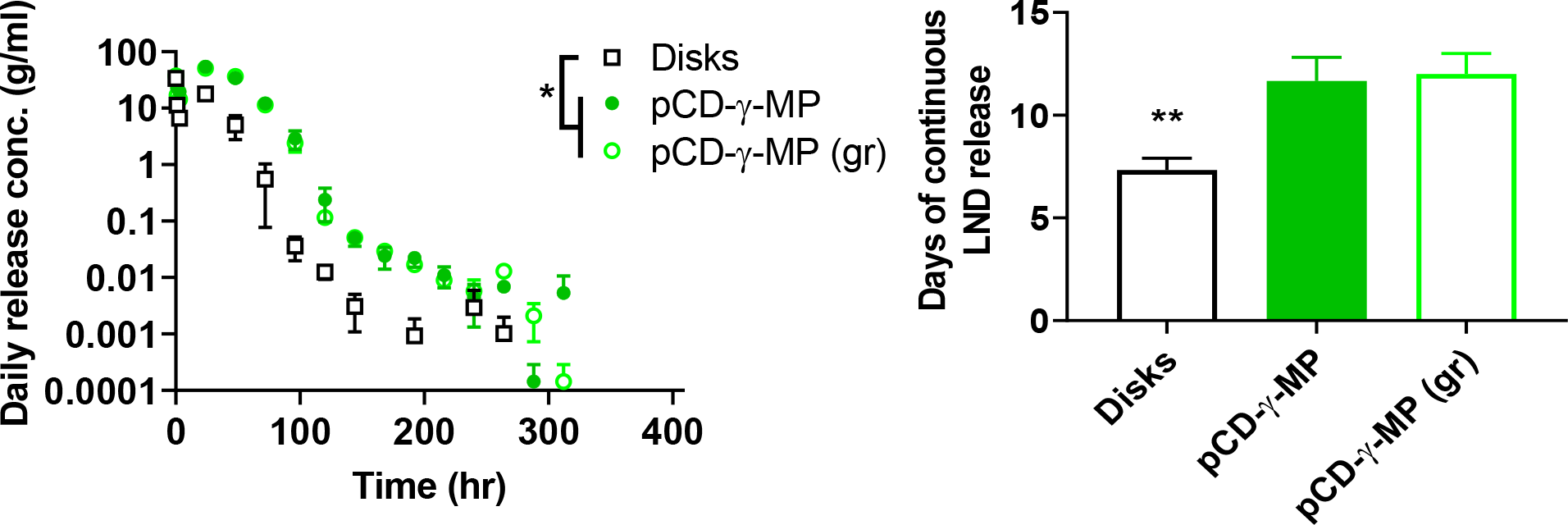
Lenalidomide daily release profiles from pCD-γ-MP, pCD-γ-MP (ground), and γ-disks (left). Days of continuous non-zero release of LND from each formulation (right). * indicates p < 0.05 by two-way ANOVA with Tukey test post-hoc between the disks and both the pCD-γ-MP and pCD-γ-ground MP groups at 3-96 hours post-incubation (left) and by one-way ANOVA with Bonferonni post-hoc for cumulative days of release between the disks and both the pCD-γ-MP and pCD-γ-ground MP (right). n=3 replicates per condition.

### 3.7 RAW 264.7 Macrophages Primarily Uptake Ground γ-CD Particles

Ground and normally synthesized pCD-γ-MP were incubated with RAW 264.7 macrophages in culture for 72 hours to assess particle uptake by cells. With the smaller size of ground particles, there was significantly increased particle uptake observed in images (Figure 7, top). Rhodamine signals from the particles were quantified and averaged per image using ImageJ (Figure 7, bottom).

**Figure 7.**
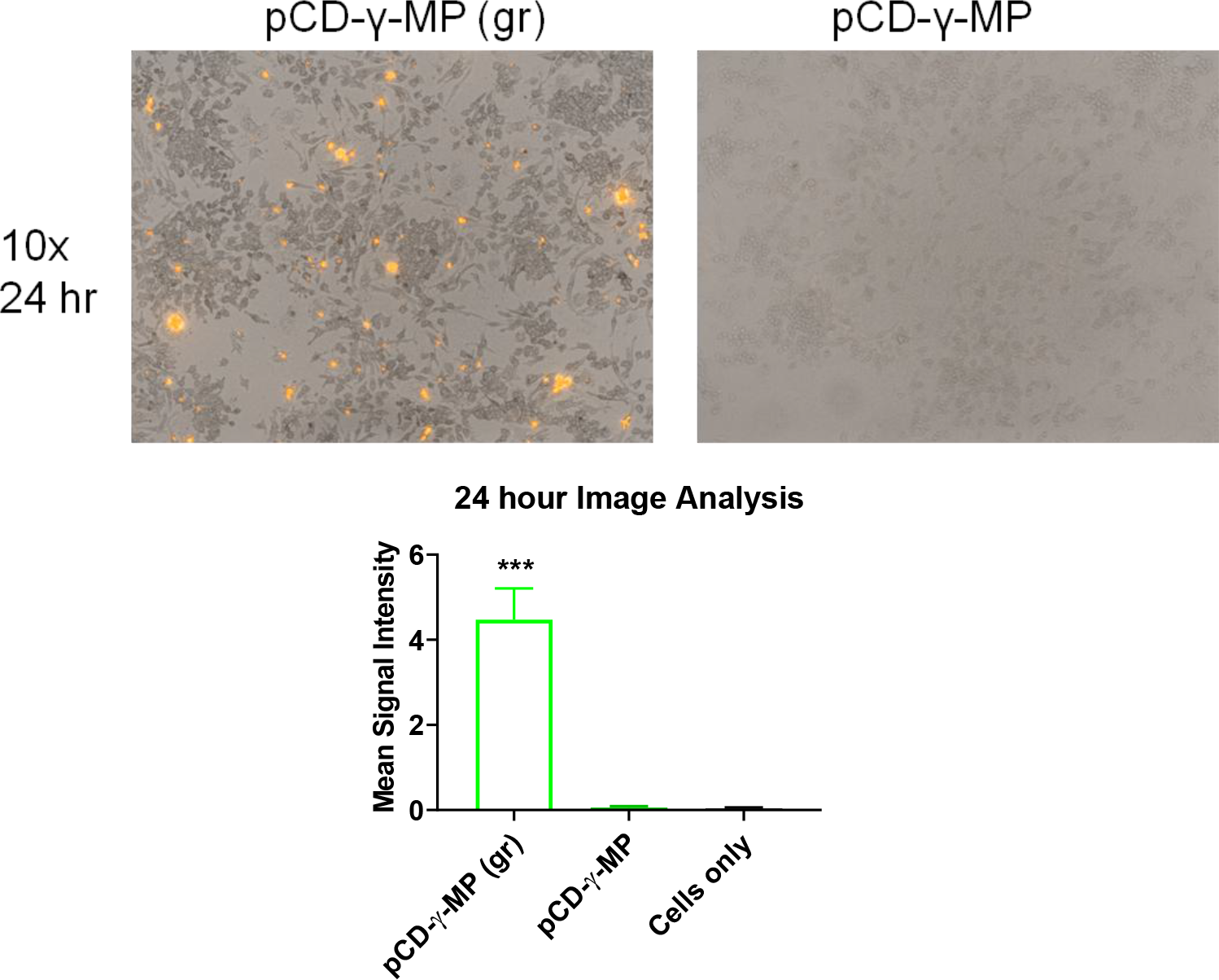
The effect of particle size on macrophage uptake *in vitro*. *** indicates p < 0.001 by one-way ANOVA with Tukey test post-hoc. n=3-7 replicates averaged from 3 images of each condition.

## 4. Discussion

We herein demonstrate the process for polymer microparticle synthesis via inverse emulsion and the effect of relevant parameters on resulting particle diameters (Figures 1 and 2). Different CD pre-polymers showed decreasing particle size for α-CD, β-CD, and γ-CD, respectively (Figure 3). Polymerized CD particles exhibit robustness to pH changes and sonication, yet particle diameter can be impacted by mortar and pestle grinding (Figure 4). Expanding these findings to research applications, we show that the monomer, but not the polymer versions of CD is cytotoxic *in vitro* (Figure 5), and correlates with previous results for CD monomer cytotoxicity ^34^. Rather than a burst of drug being delivered intravenously, CD polymer slowly releases the LND in physiological conditions for an extended period of up to 13 days (Figure 6), mostly due to the ability of γ-CD to form a drug inclusion complex of high affinity with LND, allowing it to bind into the inner pocket of CD to mediate sustained release. Further advantage of the CD polymer particle system is demonstrated in grinding particles to smaller sizes amenable to increased cellular uptake via macrophages (Figure 7). Thus these results indicate CD particle size modification can be performed independent of changing the drug release rate of the system for various applications in tissue or cellular passive targeting.

LND indicates potential for targeting numerous pathways with the greatest impacts on the polarization of macrophages, switching from M1, proinflammatory, to M2, anti-inflammatory. When the polarization of M2 macrophages is promoted, proinflammatory Th1 and Th17 cells in the peripheral lymph system and central nervous system (CNS) are subsequently inhibited. The polarization also indirectly alters proinflammatory activity of myelin-specific CD-4^+^ T cells, including IFN-γ^+^ and IL17^+^ cell subtypes ^9^. The expression levels of pro-inflammatory cytokines, such as TNF-α, interferon γ, IL-1β, and IL-6, are also reduced ^8^. LND also increases the expression and autocrine secretion of the anti-inflammatory cytokine IL-10 ^9^. Following LND treatment, the levels of anti-inflammatory cytokine IL-13 are found to be elevated ^8^. Thus far, the treatment of LND has only been delivered through oral doses, yet the benefit for selective LND localization to certain tissues and cellular subtypes is unknown. Future applications could include co-delivery of LND with other drugs for increased therapeutic effect, such as daratumumab, dexamethasone ^35^, and rituximab to treat mantle-cell lymphoma or mucosa-associated lymphoid tissue (MALT) lymphoma ^36–37^.

## Acknowledgements

The authors would like to acknowledge support from NIH T32DK083251 (NAR), R01GM12147-03 (HvR), and from an NIH Research Facilities Construction Grant (C06 RR12463-01).

